# Structural and Functional Insights into HBx-Smc6 Targeting for HBV Inhibition

**DOI:** 10.1101/2025.11.11.685253

**Authors:** Tong Cheng, Jinhong Zhou, Wenjun Huang, Lili Du, Pengyu He, Renzheng Yang, Yan Gao, Menghan Hao, Kongying Hu, Jieliang Chen, Hongxia Wang, Zihe Rao, Zhenghong Yuan, Lanfeng Wang

## Abstract

The hepatitis B virus (HBV) X protein (HBx) is a multifunctional regulator essential for HBV replication and HBV-associated hepatocarcinogenesis. Despite its central role, the molecular basis of HBx function has remained elusive. Here, we present the first cryo-electron microscopy structure of the human HBx-CRL4-Smc5/6 complex at 3.1 Å resolution. In this reconstituted ten-subunit assembly, HBx adopts a Zn^2^ □-stabilized Y-shaped architecture that simultaneously engages the DDB1 and the Smc6 subunit. A composite helix-turn-helix (HTH) pocket in HBx accommodates a conserved “Leucine Key” motif (*LRCKL*) on Smc6, forming a critical interface essential for complex stability and function. Molecular docking and biochemical validation reveal that the compound Tranilast binds this HTH pocket, disrupts the HBx-Smc6 interaction, and suppresses HBV replication. These findings define the structural mechanism by which HBx counteracts host restriction and establish the HBx-Smc6 interface as a previously unrecognized and druggable target for antiviral intervention.

## Introduction

Hepatitis B virus (HBV) infection remains a major global health challenge, affecting over 250 million people worldwide. It is a leading cause of hepatitis, cirrhosis, and hepatocellular carcinoma (HCC)^1^, accounting for nearly half of all liver cancer deaths and approximately 850,000 fatalities each year^2^. Despite effective vaccination and potent antiviral therapy, complete viral clearance is rare because covalently closed circular DNA (cccDNA) persists in infected hepatocytes^3^. The HBV X protein (HBx) is a multifunctional regulator essential for viral replication^4^ and for the transition from chronic infection to HCC^1^. HBx is a mammalian-specific innovation—viruses infecting birds or reptiles lack the X open reading frame^3^—suggesting that its emergence accompanied the evolution of persistent infection and oncogenesis in mammals. Although HBx (∼17 kDa) is highly conserved across modern and even ancient HBV strains^3^, its intrinsic disorder and conformational flexibility have long hindered structural characterization. These properties allow HBx to engage diverse host factors and to relieve cccDNA transcriptional repression^4^. Fragment-based structural studies have provided only partial insights^5^, leaving the architecture of full-length HBx undefined.

The structural maintenance of chromosomes 5/6 (Smc5/6) complex acts as a host restriction factor that suppresses HBV transcription^4^. HBx counteracts this antiviral barrier by recruiting the host Cullin-RING ligase 4 (CRL4; DDB1-Cul4A-RBx1) to ubiquitinate and degrade Smc5/6, thereby reactivating viral transcription^4^. This strategy mirrors the mechanisms employed by other viruses such as HIV, HPV, HCMV, and adenovirus, which hijack host ubiquitination machinery to inactivate restriction factors^6^. Beyond its antiviral role, Smc5/6 is essential for genome stability and DNA repair^7^, and its depletion leads to chromosomal instability and oncogenic transformation^8^. Understanding how HBx dismantles Smc5/6 is therefore key to elucidating HBV pathogenesis and identifying new antiviral targets.

## Results

We reconstituted a ten-subunit ubiquitination assembly comprising full-length HBx, CRL4, and the human Smc5/6 complex (Smc5, Smc6, Nse1-4) (Fig. 1a). The HBx-CRL4 and Smc5/6 modules were expressed separately in Sf9 cells and combined before multi-step purification, yielding a homogeneous HBx-CRL4-Smc5/6 complex (Supplementary Fig. S1b). Cryo-electron microscopy (cryo-EM) single-particle analysis revealed two major conformations: a compact 3.1 Å map (Supplementary Fig. S2) and an elongated 7.2 Å rod-like architecture resembling the yeast Smc5/6 hexamer^9^. Focused classification and local refinement produced a composite reconstruction of the full human complex (Fig. 1b; Table S1).

**Fig 1.**
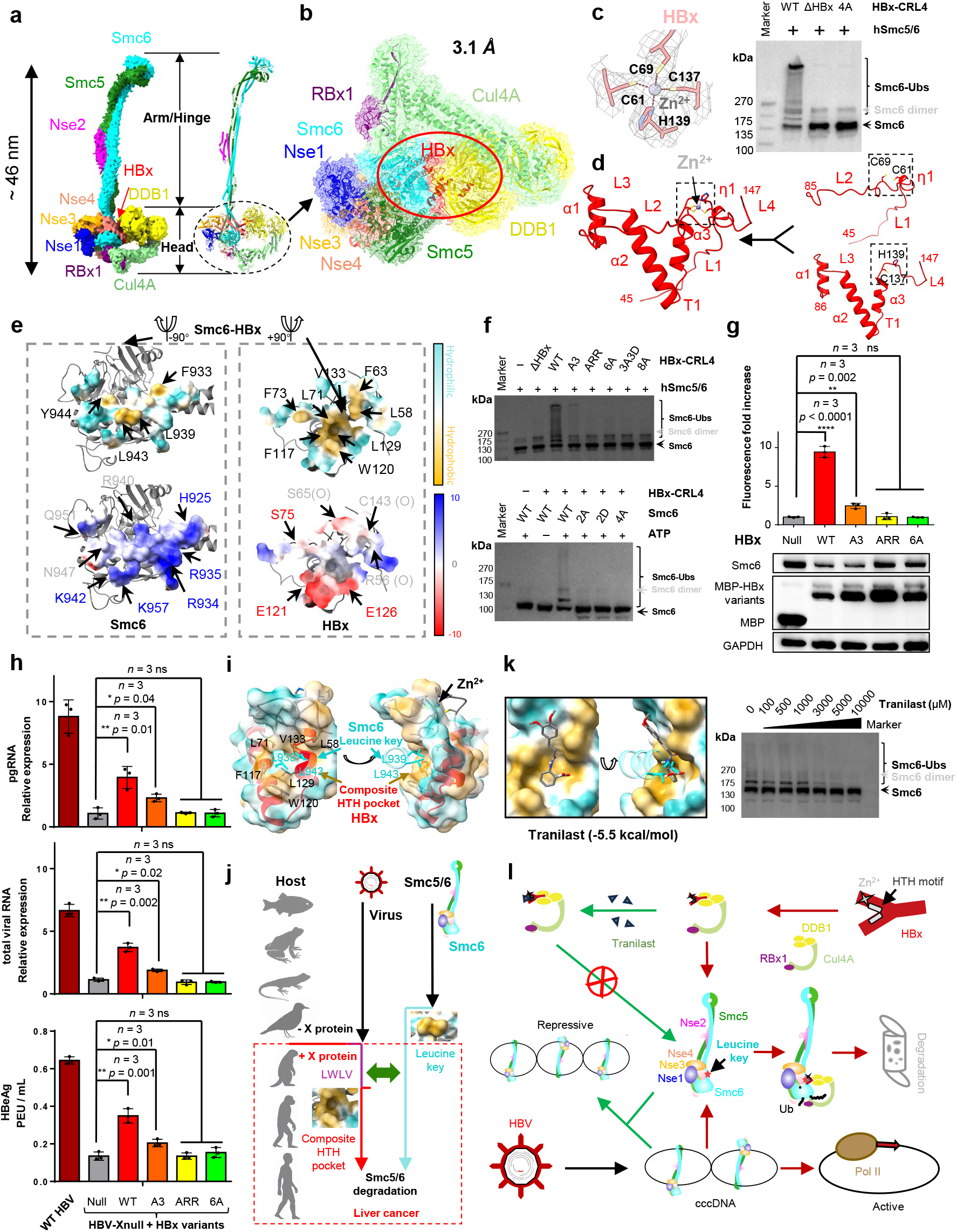
Structural and mechanistic investigation of the HBx-CRL4-Smc5/6 complex. **(a)** Cryo-EM map (left) and corresponding atomic model (right) of the full-length HBx-CRL4-Smc5/6 complex. Colors follow the scheme used throughout unless otherwise noted. **(b)** The 3.1 Å density map for the Head region with its atomic model of the HBx-CRL4-Smc5/6 complex. **(c)** Left, cryo-EM density (gray) of the Zn^2+^-binding site, tetrahedrally coordinated by C61, C69, C137, and H139 (sticks). Right, comparison of wild-type HBx and the Zn^2^ □-binding-deficient HBx-4A mutant for Smc6 ubiquitination *in vitro*. Assays were repeated at least three times with similar results. **(d)** Ribbon diagram of the HBx structure, illustrating how Zn^2+^ coordination stabilizes the interaction between the N-terminal loop and C-terminal region. **(e)** Molecular interface analysis showing surface properties of the HBx-Smc6. Top, hydrophobic contacts. Bottom, electrostatic potential. **(f)** *In vitro* ubiquitination assays. Top, wild-type HBx and its interface mutants. Bottom, wild-type Smc6 and its interface mutants. Assays were repeated at least three times with similar results. **(g)** Functional consequences of HBx mutations in HepG2 cells: Top, assessment of HBV promoter activity via the luciferase reporter assay. Bottom, determination of endogenous Smc6 protein levels by Western Blot. Data are shown as mean ± standard deviation from three independent experiments. Statistical analysis was performed using unpaired two-tailed t-tests with multiple comparisons: *p < 0.05, **p < 0.005, ****p < 0.0001. **(h)** Impact of HBx interface mutations on HBV replication in the X-null complementation system: Top, intracellular HBV pgRNA levels by RT-qPCR. Middle, intracellular total HBV RNA levels by RT-qPCR. Bottom, HBeAg secretion quantified by ELISA. Data are shown as mean ± standard deviation from three independent experiments. Statistical analysis was performed using unpaired two-tailed t-tests with multiple comparisons: *p < 0.05, **p < 0.005. **(i)** Molecular recognition mechanism: the conserved Smc6 “Leucine key” inserted into a composite HTH pocket on HBx, which is formed by the HTH motif and adjacent loops. **(j)** Co-evolutionary timeline: the emergence of the viral X open reading frame (magenta-red line), conservation of the host Smc6 “Leucine key” motif (cyan line), and the acquisition of HBV carcinogenic potential (red-dashed box, highlighting oncogenic lineages with HBx-mediated Smc5/6 degradation). **(k)** Left, Structural mechanism of Tranilast activity: (left) Predicted binding position of Tranilast (grey) within the HBx composite HTH pocket; (right) Tranilast sterically obstructs the Smc6 “Leucine key” motif to prevent productive HBx engagement. Right, dose-dependent inhibition of Smc6 ubiquitination by Tranilast *in vitro*. Assays were repeated at least three times with similar results. **(l)** Mechanistic model of HBx-mediated transcriptional regulation. HBx-induced degradation (red arrows): Zn^2^ □-stabilized HBx bridges CRL4^DDB1^ and Smc5/6^Smc6^, targeting the complex for proteasomal degradation to activate viral transcription; Tranilast inhibition (green arrows): Competitive binding to HBx prevents Smc5/6 recognition and degradation, maintaining cccDNA in a transcriptionally silent state.

In this structure, Smc5 and Smc6 form the central backbone, with Nse1-3-4 anchored at the Head and Nse2 located along the Arm/Hinge region. The HBx-CRL4 module adopts an open-ring topology: HBx binds DDB1 cleft between β-propeller domains A and C (BPA and BPC), which connect linearly to Cul4A and RBx1. Cul4A extends as a curved arm contacting Smc6, positioning RBx1 near substrate lysines for ubiquitin transfer. Thus, HBx bridges DDB1 and Smc6, forming a substrate-presentation platform optimized for Smc6 ubiquitination (Supplementary Fig. S1).

The cryo-EM density resolved HBx residues 45-147, revealing a Zn^2+^-stabilized Y-shaped structure (Supplementary Fig. S3a). The upper arms engage Smc6, whereas the lower arm inserts into the DDB1 cleft (Supplementary Fig. S1f). A tetrahedrally coordinated Zn^2+^, chelated by C61, C69, C137, and H139, stabilizes loop regions L1, η1, and L2 into a compact core (Fig. 1c and 1d). Mutation of these residues (HBx-4A: C61A / C69A / C137A / H139A) abolished Smc6 ubiquitination *in vitro* (Fig. 1c). Inductively coupled plasma-mass spectrometry confirmed stoichiometric Zn^2+^ binding to DDB1-HBx, whereas DDB1 alone lacked detectable zinc (Supplementary Fig. S3b). These results demonstrate that Zn^2+^ coordination is essential for HBx folding and function (Supplementary Fig. S3c).

At the HBx-Smc6 interface, hydrophobic and polar interactions form a stable binding surface (Fig. 1e). A hydrophobic pocket on HBx (L58, F63, L71, F73, F117, W120, L129, V133) engages a complementary region on Smc6 (Supplementary Fig. S3d). This interaction is further stabilized by hydrogen bonds and electrostatic contacts between HBx residues S75, E121, E126, and backbone atoms (R56, C143, S65) with Smc6 residues including H925, R934, R935, K942, K957, N947, Q951, and R940 (Supplementary Fig. S3e-S3g). Site-directed mutagenesis validated the functional importance of these residues: hydrophobic-core mutants (HBx-6A: L58A / L71A / F117A / W120A / L129A / V133A, HBx-8A: L58A / F63A / L71A / F73A / F117A / W120A / L129A / V133A, HBx-3A3D: L58D / L71D / F117A / W120A / L129D / V133A) and charge-disrupting variants (HBx-A3: S75A / E121A / E126A, HBx-ARR: S75A / E121R / E126R) lost or severely reduced ubiquitination activity (Fig. 1f). In HepG2 cells, HBx-WT decreased endogenous Smc6 levels, whereas HBx-6A and HBx-ARR failed to do so (Fig. 1g). HBV promoter-driven luciferase assays showed robust activation by HBx-WT, moderate by HBx-A3, and none by HBx-ARR or HBx-6A (Fig. 1g). In complementation experiments using HBV-Xnull particles, only HBx-WT restored HBeAg and pgRNA expression (Fig. 1h). Collectively, HBx residues 105-135 form a helix-turn-helix (HTH) motif that mediates Smc6 recognition (Fig. 1i; Supplementary Fig. S4a). Four of the six hydrophobic residues critical for ubiquitination (F117, W120, L129, V133) reside in this motif, while E121 and E126 form key polar contacts. Together with adjacent loop regions (L1 and L2), they create a composite HTH pocket that accommodates the Smc6 hydrophobic patch (F933, L939, L943, Y944) (Fig. 1e). Mutation of these Smc6 residues (Smc6-2A: L939A / L943A, Smc6-2D: L939D / L943D, Smc6-4A: F933A / L939A / L943A / Y944A) abolished ubiquitination (Fig. 1f). L939 and L943, extending from the conserved LRCKL motif, form the “Leucine key” that inserts into the HBx composite HTH pocket (Fig. 1i), which is the structural Achilles’ heel of Smc5/6.

The HBx-DDB1 interface buries ∼5180 Å^2^ of surface area, far exceeding that of isolated H-Box (∼1667 Å^2^) or HLH (∼1972 Å^2^) (Supplementary Fig. S5). Although Zn^2+^ is dispensable for DDB1 association^10^, it is indispensable for Smc6 engagement, indicating that metal coordination stabilizes the substrate-binding pocket rather than the adaptor interface. While previous studies have been limited to fragmentary structures of individual HBx motifs^5^, our reconstitution of the HBx-CRL4-Smc5/6 assembly enabled the determination of its three-dimensional structure in its most complete form at 3.1 Å resolution. The Zn^2+^-tethered architecture (residues 45-147) links flexible loops to the HTH motif, forming a composite pocket accommodating the Smc6 Leucine key. This compact fold differs markedly from the extended α-helix observed in Bcl-2-bound HBx, highlighting its conformational adaptability as a “macro-molecular glue” (Supplementary Fig. S6). In contrast, AlphaFold-predicted HBx models misassigned the Smc6 interface and lacked the Zn^2+^ core (Supplementary Fig. S7), emphasizing the necessity of experimental data. Our cryo-EM analysis also provides the first architectural insights into the human Smc5/6 complex. Despite low sequence identity (13-25%) with yeast orthologs, the overall ∼46 nm rod-like shape is conserved (Supplementary Fig. S8). However, the human complex exhibits distinct rearrangements: relative to the aligned Smc6 Head, the Hinge is shifted ∼86 Å, the Smc5 Head rotated ∼18 Å, and Nse4 and Nse1-3-4 are displaced by up to 40 Å and 15 Å, respectively. These features likely reflect higher-eukaryotic regulatory adaptations. The exposed Leucine key within the Head domain constitutes an intrinsic vulnerability exploited by HBx.

Functionally, HBx-mediated Smc5/6 degradation produces two synergistic outcomes: it lifts epigenetic repression of cccDNA to enable viral transcription, and it compromises host genomic stability, promoting oncogenic transformation. Because Smc5/6 is indispensable for replication-fork maintenance and DNA repair^7^, its loss leads to chromosomal instability^8^. By co-opting CRL4 to degrade Smc5/6, HBx subverts a genome-stabilizing complex to sustain infection. Evolutionary analysis revealed that the Smc6 Leucine key motif emerged in birds and became conserved in mammals (Fig. 1j; Supplementary Figs. S4b-S4c). Concomitantly, the HBx gene arose in mammalian hepadnaviruses, indicating co-evolution of viral pocket and host key. Both human and woodchuck HBx share conserved FWLV residues in the HTH motif and mediate Smc5/6 degradation^4^. This adaptation likely provided a selective advantage by allowing hepadnaviruses to bypass Smc5/6-mediated silencing, linking persistence to oncogenesis. Given the evolutionary conservation of the HBx-Smc6 interface, we explored its druggability. Tranilast, previously reported to inhibit HBx-dependent replication^11^, suppressed Smc6 ubiquitination by the HBx-CRL4 complex in a dose-dependent manner (Fig. 1k). Molecular docking showed that Tranilast occupies the HTH pocket, sterically blocking the Smc6 Leucine key (Fig. 1k). Across all HBV genotypes, the hydrophobic and charged residues of the HTH pocket and Zn^2+^ cluster are invariant (Supplementary Fig. S9) and preserved in ancient HBx sequences^3^, reflecting extraordinary stability. Unlike rapidly mutating viral surface proteins^12^, the HBx core has remained unchanged for millennia. Clinically validated protein-protein interaction inhibitors^13^ demonstrate the feasibility of targeting such interfaces. Currently, no approved antiviral drug directly targets HBx and existing nucleos(t)ide analogs suppress replication but fail to eradicate cccDNA^14^. By stabilizing Smc5/6 and restoring its antiviral function, small molecules like Tranilast could provide a complementary therapeutic route toward functional cure (Fig. 1l).

In summary, we resolved the near-atomic architecture of the ten-subunit HBx-CRL4-Smc5/6 complex and defined the structural mechanism by which HBx hijacks the host CRL4 ligase to degrade Smc5/6. HBx adopts a Zn^2+^-stabilized Y-shaped conformation that bridges DDB1 and Smc6 through a composite HTH pocket accommodating the Smc6 Leucine key. Mutation or small-molecule occupation of this pocket abolishes Smc6 ubiquitination and suppresses HBV replication. This evolutionarily conserved interface represents a mutation-resistant and chemically tractable target for antiviral drug development. By revealing a molecular Achilles’ heel in the host restriction machinery, our work provides a structural foundation for next-generation therapeutics designed not merely to suppress HBV but to restore the host’s intrinsic antiviral defense.

## Data availability

The cryo-EM maps and atomic coordinates have been deposited into the Electron Microscopy Data Bank and the Protein Data Bank with the following accession codes: EMD-63446 (PDB: 9LWI, the Head region of the HBx-CRL4-Smc5/6 complex), EMD-63447 (PDB: 9LWJ, the Head/Arm region of the HBx-CRL4-Smc5/6 complex), EMD-63448 (PDB: 9LWK, the Hinge/Arm region of the HBx-CRL4-Smc5/6 complex), and EMD-63449 (PDB: 9LWL, the HBx-CRL4-Smc5/6 complex).. Structures used for structural analysis have the following accession codes from the Protein Data Bank: 3I7H, 2HYE, 5WY5, 8I13, 8GTX, 8WLS, 5FCG, 5B1Z, 9J6K. The AlphaFold identifiers used for model building include AF-Q8IY18-F1 (hSmc5), AF-Q96SB8-F1 (hSmc6), AF-Q96MF7-F1(NSMCE2/Nse2), AF-Q96MG7-F1 (NDNL2/Nse3), AF-Q8N140-F1 (EID3/Nse4). Accession numbers are listed in the key resources table. Any additional information required to reanalyze the data reported in this paper is available from the lead contact upon request.

## Acknowledgements

We thank all the staff members of the cryo-EM facilities at Shanghai Institute of Immunity and Infection, CAS and Fudan University (Fenglin Campus) for their assistance with cryo-EM data collections. This work was supported by grants from the National Key R&D Program of China (2022YFA1303600, to L.W.), the Strategic Priority Research Program of the Chinese Academy of Sciences (XDB29010205, to L.W.), and the Shanghai Municipal Science and Technology Major Project (2019SHZDZX02, to L.W.).

## Author contributions

L.W., conceived, Z.Y., and Z.R. coordinated this project. T.C., J.Z., and L.D. carried out molecular cloning, cell culture, and protein preparation. T.C. collected EM data with the assistance of G.Y.. L.W. and T.C. built structural models. P.H. carried out molecular docking. T.C. and J.Z. conducted *in vitro* ubiquitination assay. J.Z., L.D., R.Y., and H.W. carried out the cellular experiments. W.H., M.H., K.H., J.C., and Z.Y. carried out HBV-associated experiments. L.W., Z.Y., and Z.R. analyzed the results. L.W., T.C., and J.Z. wrote the manuscript with input from all the authors.

