## Supplementary information, Figures and Table for "Structural and Functional Insights into HBx-Smc6 Targeting for HBV Inhibition"

**Supplementary information, Table S1. Cryo-EM data collection, refinement, and validation statistics for HBx-CRL4-Smc5/6 complex**

|  | Head  EMDB-63466  PDB-9LWI | Head/Arm  EMDB-63447  PDB-9LWJ | Hinge/Arm  EMDB-63448  PDB-9LWK | Composite  EMDB-63449  PDB-9LWL |
| --- | --- | --- | --- | --- |
| **Data collection and processing** |  |  |  |  |
| Magnification | 130,000 | 130,000 | 130,000 | 130,000 |
| Voltage (kV) | 300 | 300 | 300 | 300 |
| Electron exposure (e–/Å^2^) | 48.5 | 48.5 | 48.5 | 48.5 |
| Defocus range (μm) | -1.2 to -2.2 | -1.2 to -2.2 | -1.2 to -2.2 | -1.2 to -2.2 |
| Voxel size (Å) | 0.932 | 1.864 | 1.864 | 1.864 |
| Symmetry imposed | C1 | C1 | C1 | C1 |
| Initial particle images (no.) | 533,889 | 69,510 | 69,510 | 69,510 |
| Final particle images (no.) | 401,828 | 20,327 | 29,615 | 69,510 |
| Map resolution (Å)  FSC threshold | 3.12  0.143 | 7.19  0.143 | 7.32  0.143 | 7.25  0.143 |
| Map resolution range (Å) | 7-3 | 9-5 | 9-5 | 9-5 |
| **Refinement** |  |  |  |  |
| Initial model used (PDB code) | n/a | n/a | n/a | n/a |
| Model resolution (Å)  FSC threshold | 3.1  0.143 | 7.1  0.143 | 7.1  0.143 | 7.1  0.143 |
| Model resolution range (Å) | 7-3 | 9-5 | 9-5 | 9-5 |
| Map sharpening *B* factor (Å^2^) | -121.9 | -521.0 | -587.5 | n/a |
| Model composition  Non-hydrogen atoms  Protein residues  Ligands | 28126  3502  Zn:6 | 33723  4185  Zn:6 | 6084  736  0 | 39738  4914  Zn:6 |
| *B* factors (Å^2^)  Protein  Ligand | 41.46  75.96 | 361.25  573.19 | 362.47  --- | 359.96  573.19 |
| R.m.s. deviations  Bond lengths (Å)  Bond angles (°) | 0.007  1.100 | 0.004  0.881 | 0.004  0.891 | 0.004  0.896 |
| Validation  MolProbity score  Clashscore  Poor rotamers (%) | 1.66  8.08  0.64 | 1.60  9.25  0.37 | 1.47  8.82  0.59 | 1.63  9.53  0.41 |
| Ramachandran plot  Favored (%)  Allowed (%)  Disallowed (%) | 96.60  3.40  0.00 | 97.47  2.53  0.00 | 98.09  1.91  0.00 | 97.34  2.66  0.00 |

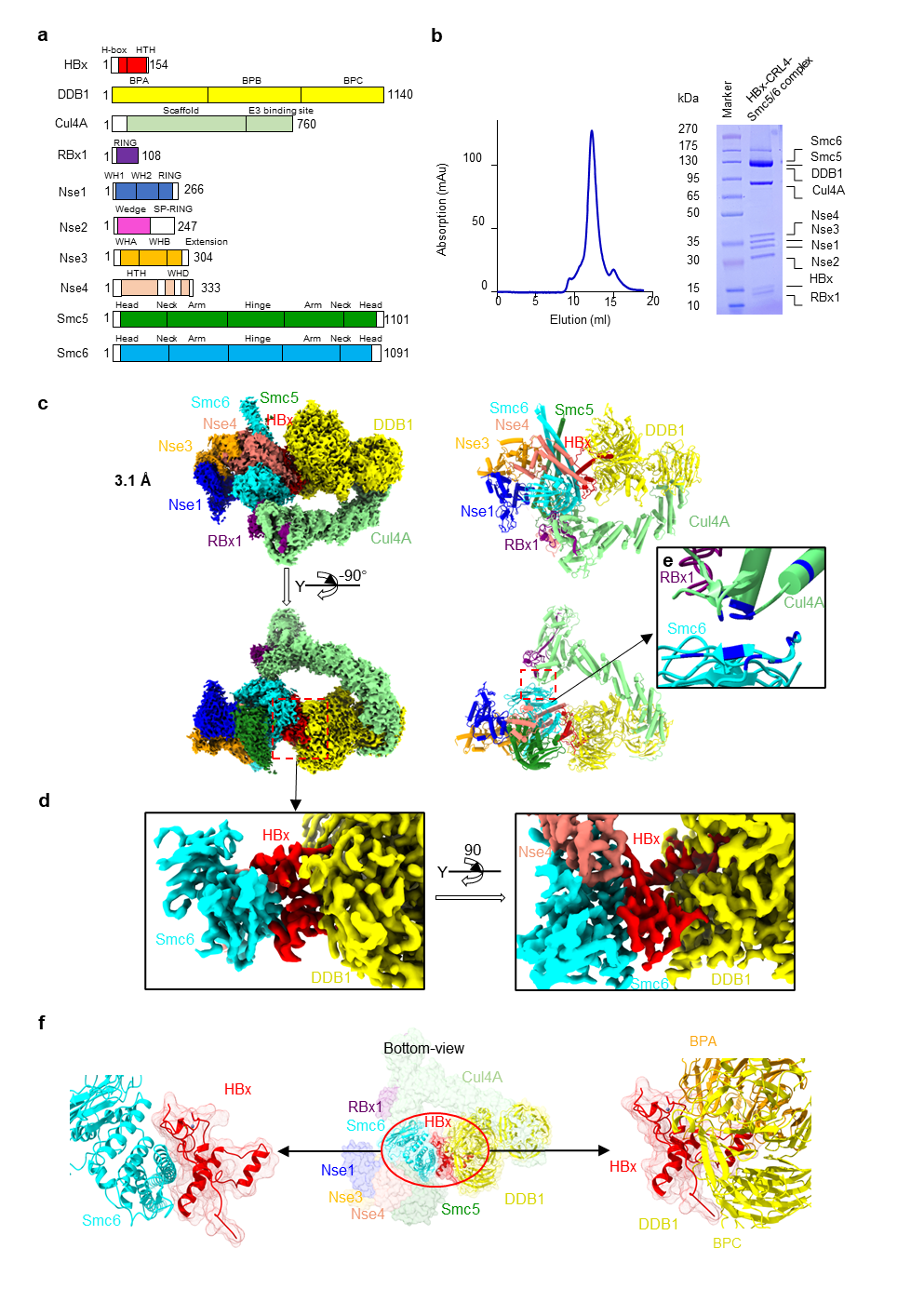

**Supplementary information, Fig. S1. Structure of the HBx-CRL4-Smc5/6 complex.**

(**a**) Domain organization of the ten subunits in the HBx-CRL4-Smc5/6 complex. Resolved structural regions from this study are colored.

(**b**) Size-exclusion chromatography elution profile of the HBx-CRL4-Smc5/6 complex, with Coomassie blue R250-stained SDS-PAGE of peak fractions. Purification was independently repeated at least three times with consistent results.

(**c-e**) High-resolution structural features of the complex: (**c**) Cryo-EM density (left) and model (right) of the Head region; (**d**) Detailed interface between Smc6, HBx, and DDB1; (**e**) Cul4A-Smc6 interaction site (highlighted in blue).

(**f**) Surface representation of HBx (red) bridging DDB1 (yellow) and Smc6 (cyan), viewed from the bottom.

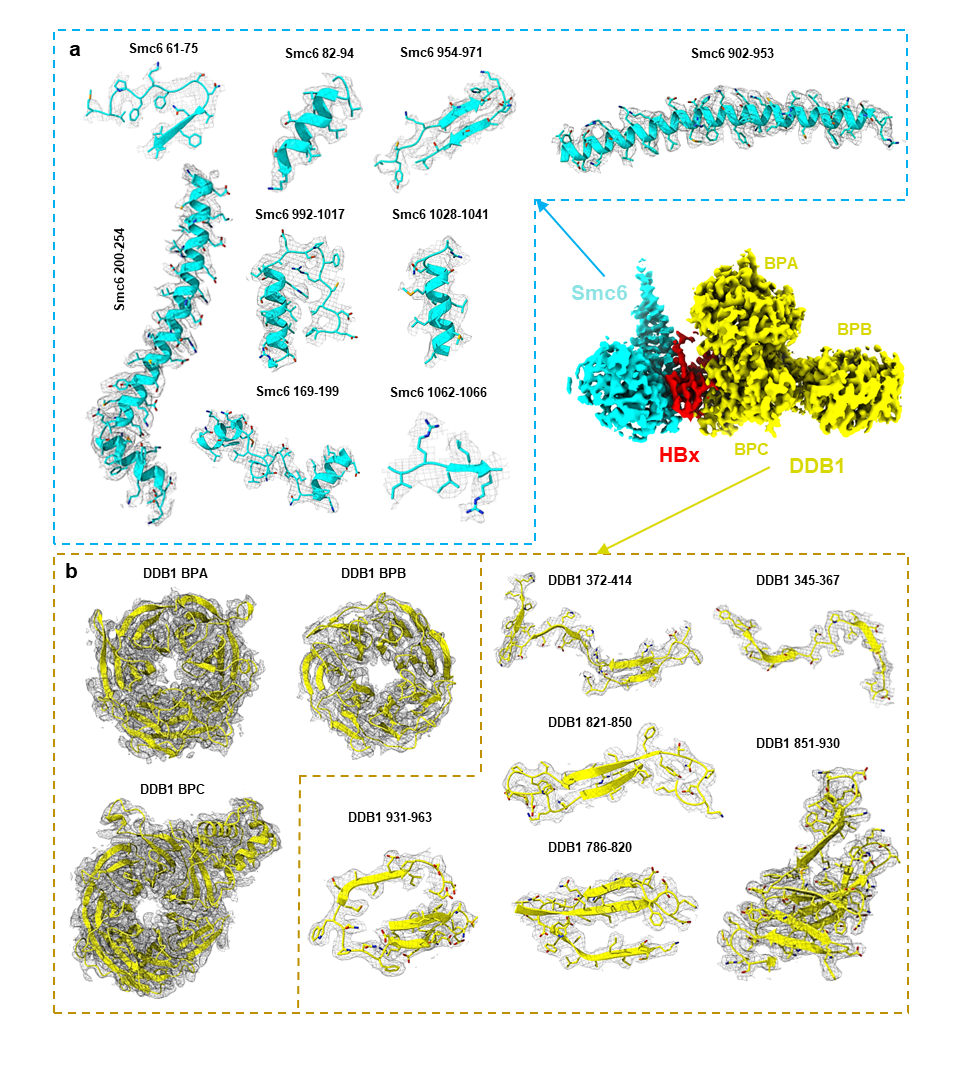

**Supplementary information, Fig. S2. Local density maps of Smc6 and DDB1**

(**a** and **b**) High-resolution density maps of Smc6 (**a**) and DDB1 (**b**) from the HBx-CRL4-Smc5/6 complex.

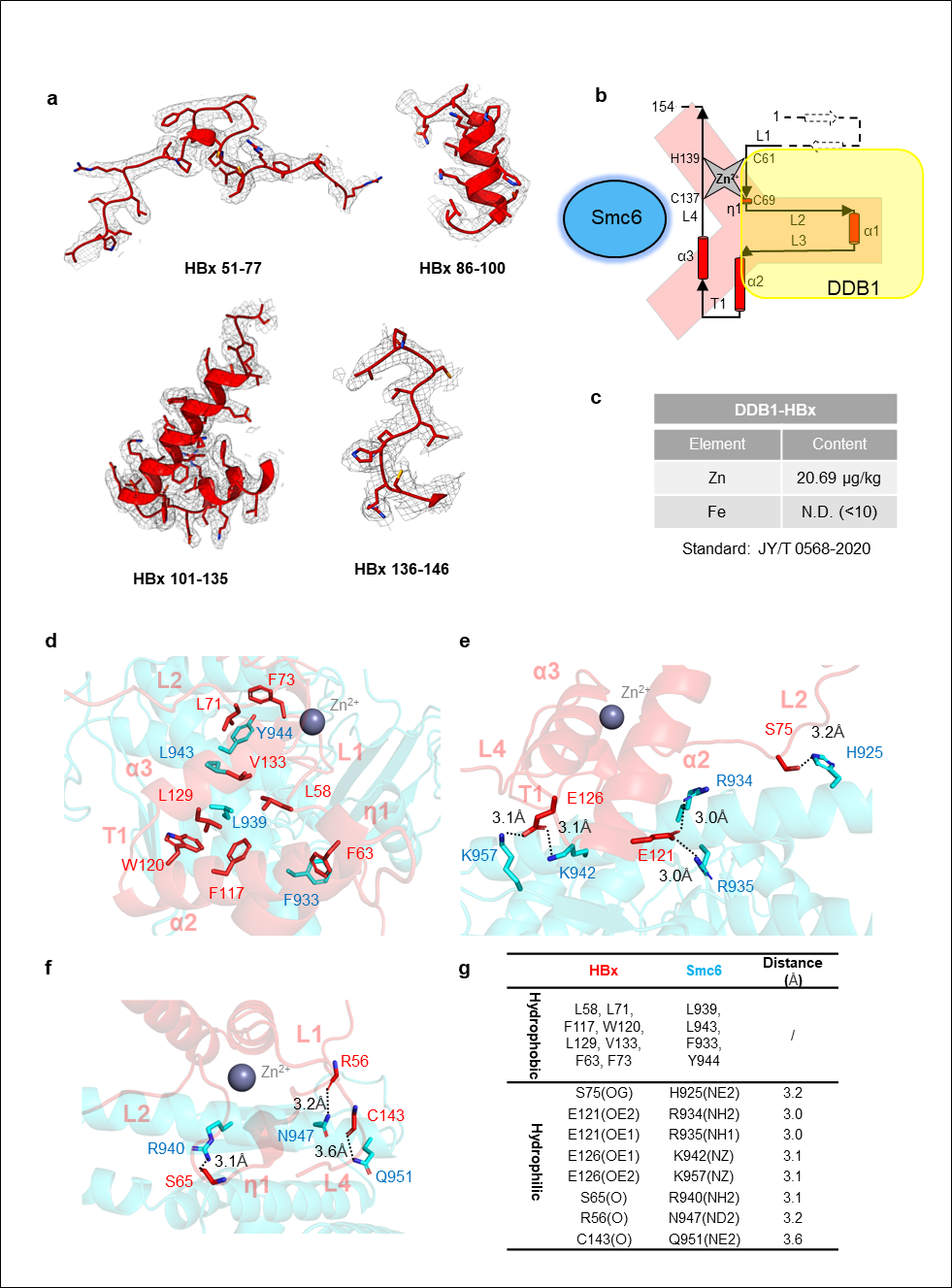

**Supplementary information, Fig. S3. Local density and HBx-Smc6 interactions.**

(**a**) High-resolution local density maps of HBx.

(**b**) Topological diagram of HBx illustrating its Zn^2+^-stabilized Y-shaped architecture that bridges DDB1 and Smc6.

(**c**) ICP-MS analysis confirming Zn^2+^ stoichiometry in the DDB1-HBx complex.

(**d-f**) Stick representations of hydrophobic (**d**) and hydrophilic (**e** and **f**) interactions between HBx and Smc6, including side-chain (**e**) and main-chain (**f**) contributions.

(**g**) List of key HBx residues, with the corresponding hydrogen bond distances explicitly indicated.

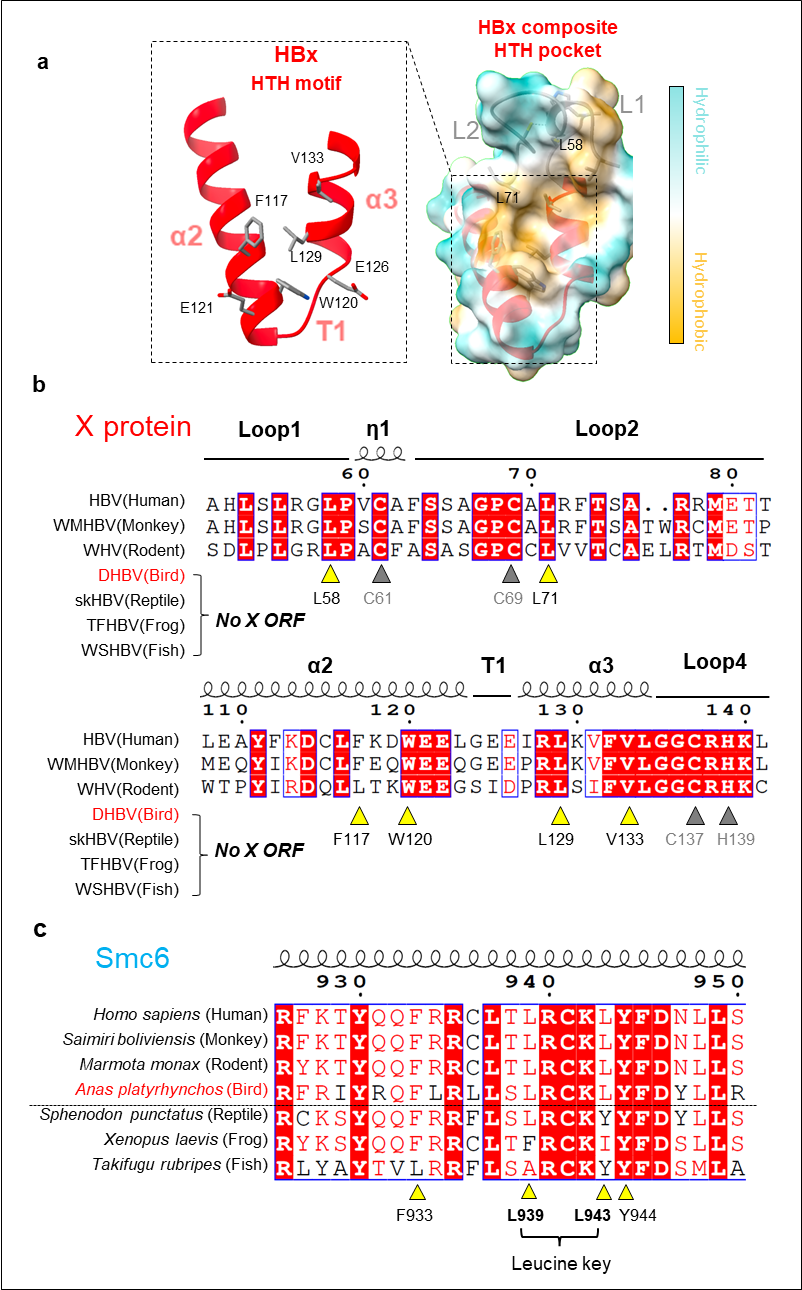

**Supplementary information, Fig. S4. Conservation analysis of Smc6-HBx interaction key residues.**

(**a**) Structural model of HBx novel helix-turn-helix (HTH) motif (α2-T1-α3, left), and HBx composite HTH pocket (right).

(**b**) Hepadnaviral X-protein alignment highlighting: Composite HTH pocket residues (yellow triangles) and Zn²⁺-chelating sites (gray triangles).

(**c**) Evolutionary conservation of Smc6 interface residues across vertebrates (bottom-to-top: increasing complexity), with Leucine key highlighted.

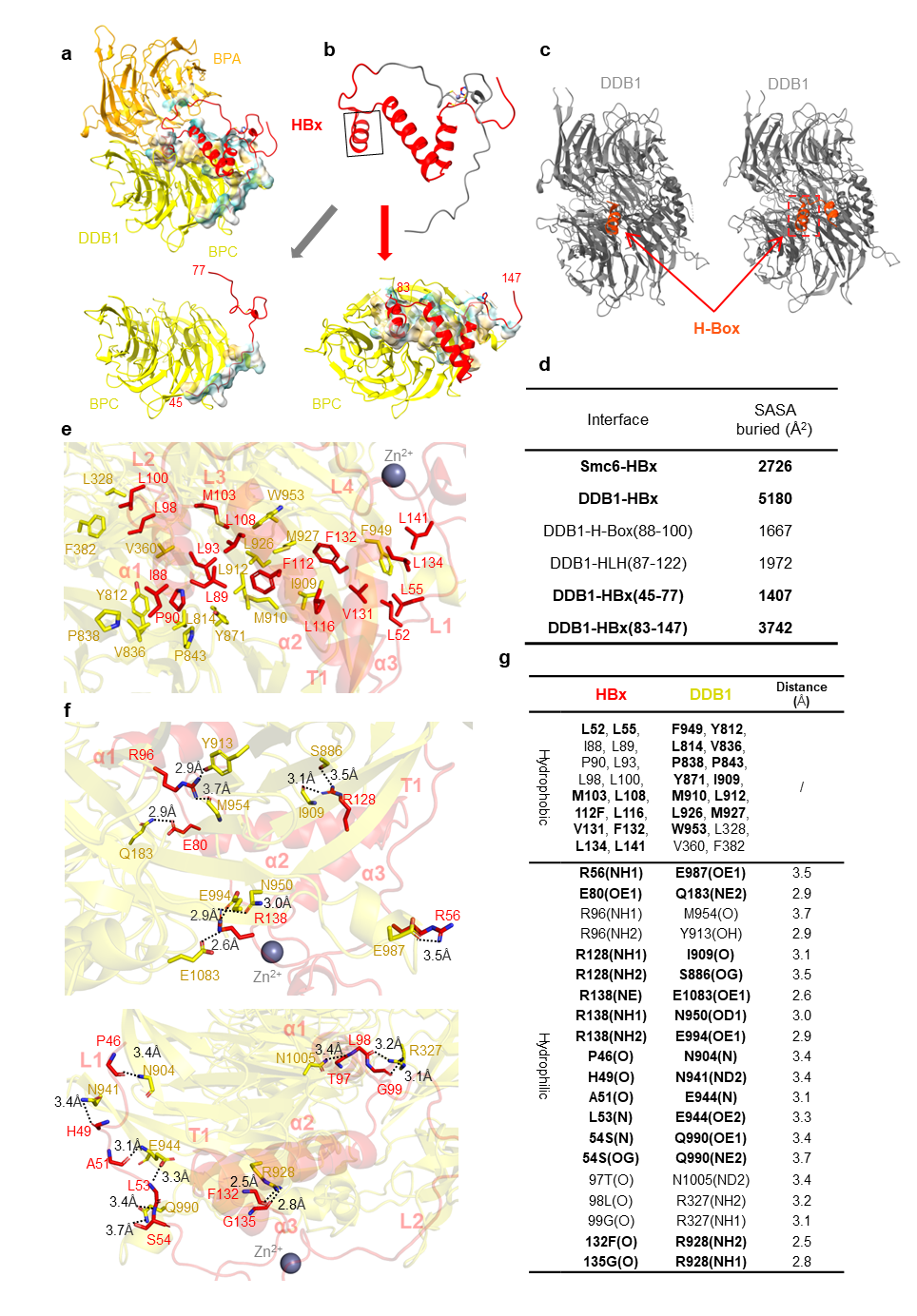

**Supplementary information, Fig. S5. Structural analysis of HBx-DDB1 interactions.**

(**a**) Cartoon representation of the HBx interacting with DDB1 (majority in BPC domain).

(**b**) Detailed view of HBx-DDB1 binding interfaces: (bottom left) N-terminal HBx interactions (residues 45-77); (bottom right) C-terminal HBx interactions (residues 83-147).

(**c**) Structural comparison of the H-Box motif (PDB: 3I7H)^1^ and HLH motif (PDB: 9J6K)^2^ interacting with DDB1 in the previous structures.

(**d**) Comparison of the buried surface areas of six HBx-related interfaces.

(**e-g**) Interactions between HBx and DDB1: (**e**) Stick representation of the hydrophobic interactions; (**f**) hydrogen bond interactions; (**g**) the key residues and the hydrogen bond distances were listed, with the newly identified interactions beyond H-Box in bold.

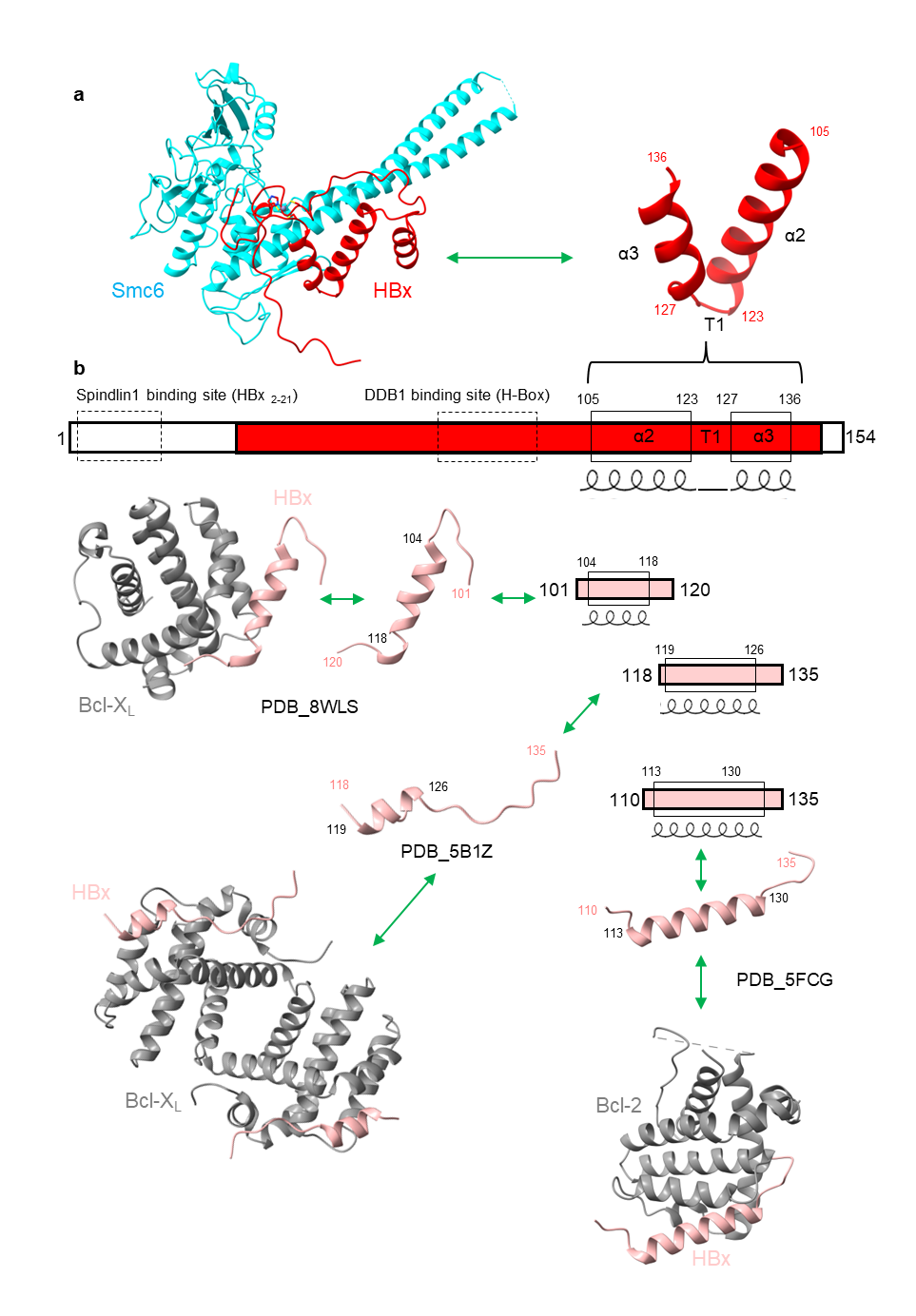

**Supplementary information, Fig. S6.** **Structural plasticity of HBx with host factors.**

(**a**) Cartoon representation of the HTH motif from this study.

(**b**) Comparison of our structure with previous HBx fragments binding Spindlin1 (ref. ^3^), DDB1 (ref. ^1,2^), Bcl-2 (PDB: 5FCG)^4^ and Bcl-X_L_ (PDBs: 8WLS^5^ and 5B1Z^6^).

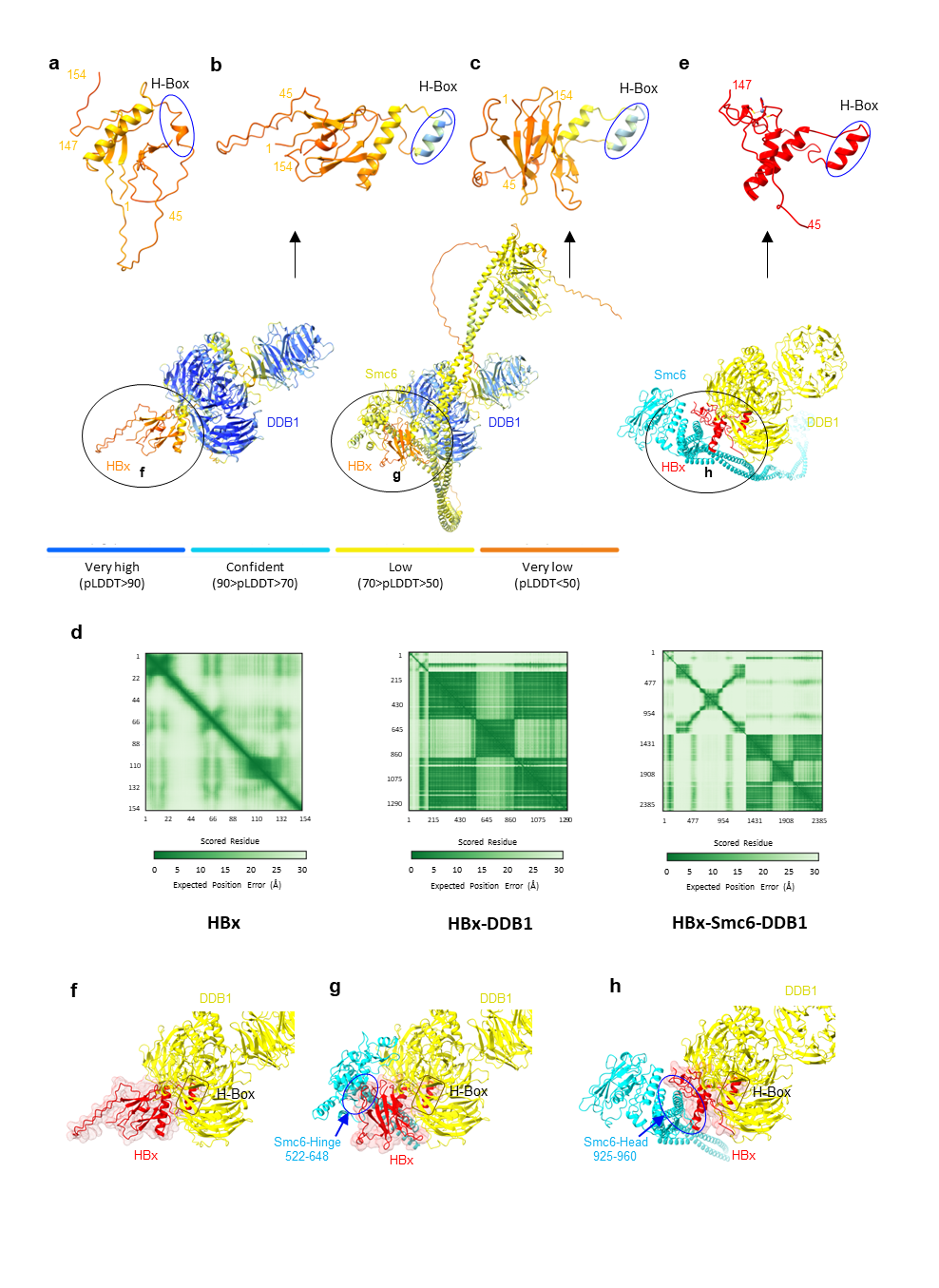

**Supplementary information, Fig. S7. Comparison of experimental and AlphaFold3-predicted HBx models.**

(**a-d**) AlphaFold3-predicted models of apo HBx (**a**), HBx-DDB1 (**b**), and HBx-Smc6-DDB1 (**c**), with model scores and PAE plot (**d**).

(**e**) Experimental HBx structure from this study.

(**f-h**) Interface comparison between predicted (**f** and **g**) and experimental (**h**) structures.

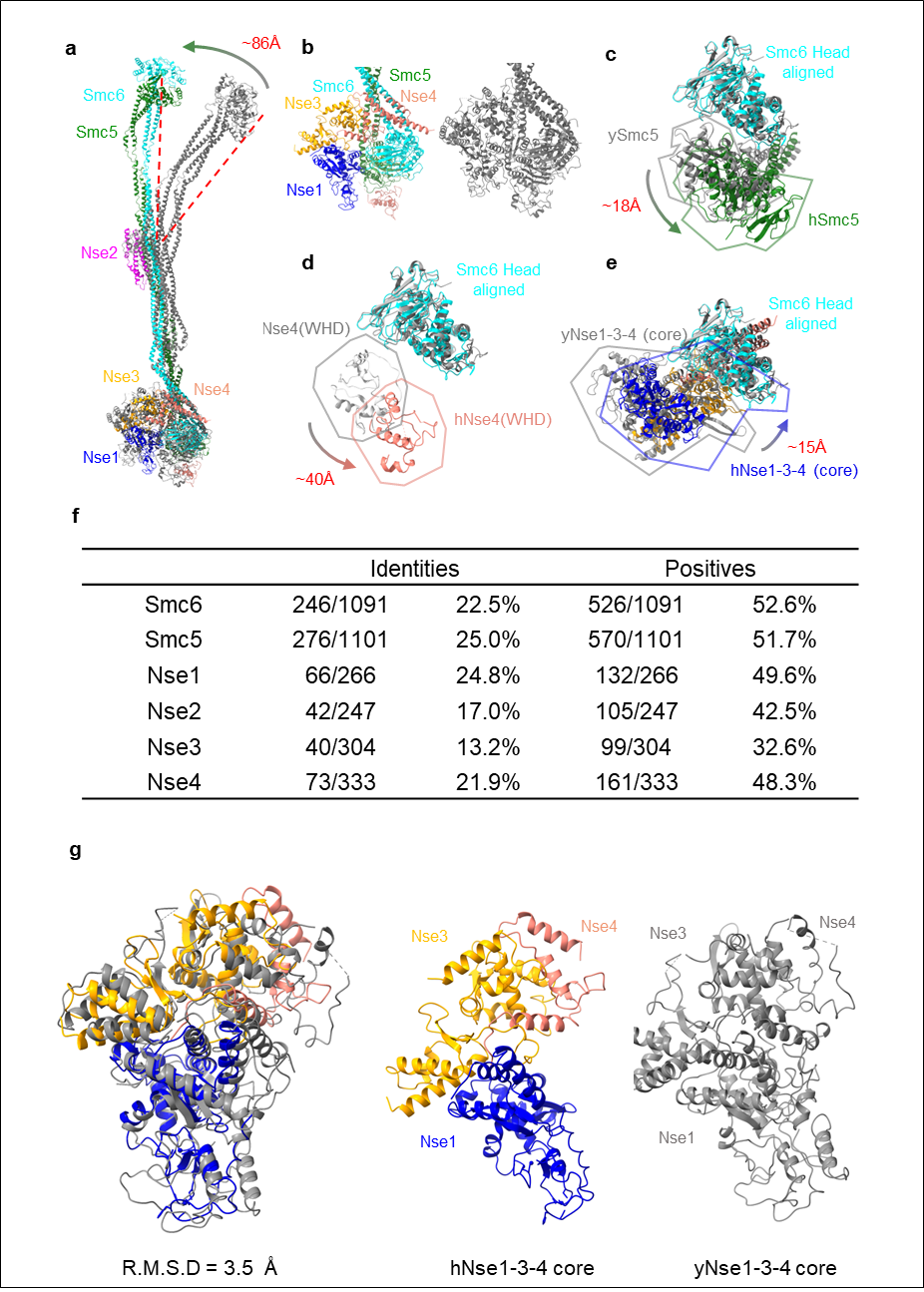

**Supplementary information, Fig. S8.** **Sequence and structure comparison of the human and yeast Smc5/6 complexes.**

(**a-e**) Structural superimposition of hSmc5/6 (colored) and ySmc5/6-6mer (gray; PDB: 8I13), aligned at the Smc6 Head domain. Highlighted regions include: (**b**) overall Head regions, (**c**) Smc5 Head, (**d**) Nse4 C-terminal WHD, and (**e**) Nse1-3-4 core.

(**f**) Sequence conservation of the six subunits in human and yeast Smc5/6 complexes.

(**g**) Structural comparison of Nse1-3-4 core from hSmc5/6-6mer (color) and ySmc5/6-6mer (PDB: 8WJN, grey).

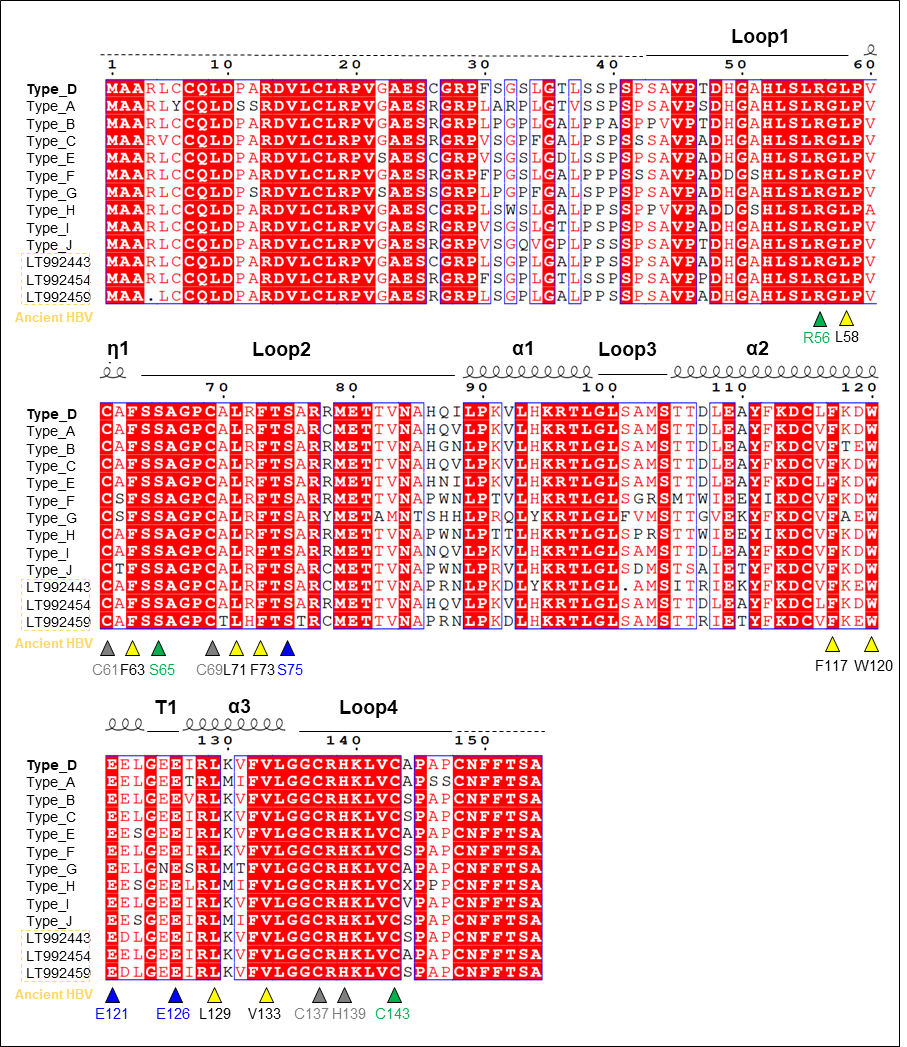

**Supplementary information, Fig. S9. Conserved key HBx residues across HBV genotypes and ancient strains.**

Sequence alignment of HBx variants from ten HBV genotypes and ancient HBV sequences. Conserved residues involved in the Smc6 interface are highlighted: hydrophobic (yellow triangles), side-chain polar/charged (blue triangles), backbone (green), and Zn^2+^-chelating (gray triangles).
